# Molecular Dynamics Study of Dehydrated Lipopolysaccharide Membrane

**DOI:** 10.1101/2020.05.31.126326

**Authors:** Changjiang Liu, Paolo Elvati, Angela Violi

## Abstract

The outer membrane of bacteria is known to play an important role in the rapid response to desiccation, although the causes and the extent of these effects are still mostly unclear. For this reason, in this work we study the desiccation response of the Gram-negative lipopolysaccharide (LPS) bacterial outer membranes. By analyzing molecular dynamics simulations of LPS membranes of different composition during desiccation, we identified the formation of a rigid protective layer of polysaccharides that not only reduces the water evaporation but is also able to indirectly preserve several structural features and as such membrane functionality. Overall, we found that the presence of polysaccharide layer is critical in the conservation of a layer of water in proximity of the hydrophobic region as well as the structure of the lipids acyl chain structure.

## Introduction

Desiccated environment is harmful for microbial survival. This fact provides an efficient approach to kill pathogenic bacteria or suppress their growth with many applications, such as food preservation, antimicrobial and disinfection on surface. However, it also becomes a severe problem for the agricultural industry when the survival of some beneficial soil bacteria is affected by droughts.

The natural resistance of bacteria to desiccation has been extensively studied. Structurally, several polymeric layers have been identified on the surface of bacterial cell of being responsible for the bacterial resistance to desiccation, such as the outer membrane of lipopolysaccharide (LPS) adapted by Gram-negative bacteria,^1,2^ the peptidoglycan layer by Gram-positive bacteria,^3,4^ and extracellular matrices common in both.^5–7^ Meanwhile, bacteria adapt several biosynthetic responses to environmental desiccation such as formation of biofilms, formation of spores, dormancy, and gene-regulated synthesis of anti-desiccation materials.^8^ These two types of resistance are often complementary to each other: as the biosynthetic pathways are slow in response, the polymeric layers are immediately responsible for protecting the cell when facing desiccation. Once bacteria survived the initial desiccation, they activate specific biosynthetic pathways, including some pathways which synthesize the units for thickening the polymer layers. ^9–13^

While there has been a lot of progress in the identification of the chemical components of these layers, as well as the bio-synthetic pathways and gene regulations, the structure and functions of these polymer layers has not been fully understood due to their complexity. For example, even considering only LPS molecules, there is a great variety in the type of repeating units (more than 190 types being identified for the *E. coli*), length (usually from 0 to 50 units) and connectivity (linear or branched).^1^ Moreover, how such structures are organized in space, how the structures change in response to the environmental desiccation, and what the mechanism is of such structural change being associated with bacterial resistance to desiccation are still open questions.

Due to the complexity of these systems, molecular models have been successfully employed to study the key characteristics of these membranes, with molecular dynamics simulations showingXxxxx to be able to match experimental measures like atomic distances, order parameter, lipid packing, and area per lipid.^14–16^ Despite this successes, these computational techniques have note been applied to understanding the mechanisms of the short term dehydration resistance observed for LPS membranes.

In this work, we fill this gap by studying the effect of desiccation on LPS membranes. Specifically, we are interested in understanding how the structure of the outer membrane changes and how such changes affect its functionality when only limited amount of water is present. By comparing the LPS membranes at different hydration level, we identified several significant structural changes that alter LPS membrane during the desiccation that result in the formation of a polysaccharide layer that reduces the evaporation of the water close to the membrane head groups. The simulations show how many changes and the preservation of the membrane functionality is strongly affected by the presence of polysaccharide layer, which seems to be critical in the short term response to desiccation.T

## Methodology

### MD simulations

We used NAMD^17^ version 2.13 with CHARMM force field verwith a timestep of 2 fs while keeping all the C– H and O – H bonds rigid via the SHAKE algorithm.^20^ Long range electrostatic interactions were modeled with the particle mesh Ewald method^21^ using a 0.1 nm grid spacing, a tolerance of 10^−6^ and cubic interpolation. Temperature was kept constant at 310 K using a Langevin thermostat^22^ with a time constant of 2 ps. Non-bonded short-range interactions where smoothly switched to 0 between 1 and 1.2 nm with a X-PLOR switch function.

System were simulated with cubic periodic boundary conditions in either canonical ensemble or NPsT ensemble. In the latter, the x and y dimensions, which correspond to the plan parallel to the bilayers, are coupled but allowed to change independently from the z dimension, the membrane normal. For NPsT ensemble, the Nosé-Hoover Langevin piston method^23,24^ with a period of 50 fs a and decay of 25 fs was applied to keep the average pressure at 101.325 kPa.

### System preparation

We prepared two different bilayers, both with a leaflet consisting of 48 1,2-dipalmitoyl-sn-glycero-3phosphoethanolamines (DPPEs) (inner leaflet) while the other was composed by 16 LPSs with a different number of repeating units (outer leaflet). The topology of the LPS molecule in both bilayers was generated using the CHARMM-GUI LPS Modeler^25,26^ as *E. coli* R1 (core) O6 (antigen) molecules. The difference between the systems is limited to the number of the repeating units in the LPS: 5 for the DPPE/LPS-5ru system (see Fig. 1A and Fig. S1 for more details on the LPS structure) and 0 for the DPPE/LPS-0ru system.

**Figure 1:**
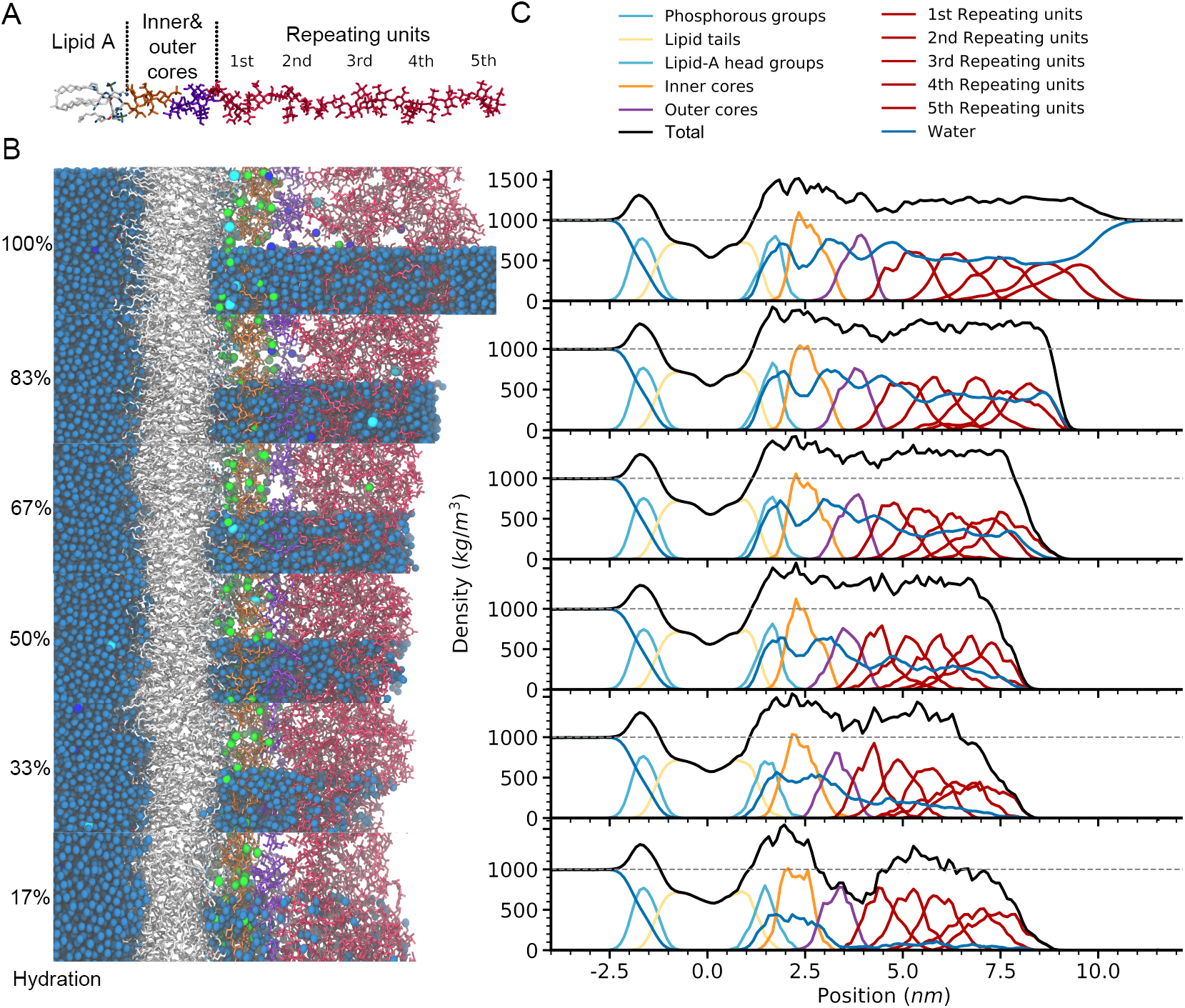
LPS membrane structure and density distribution. LPS (panel A) can be divided in four regions: lipid A, inner core, outer core and a variable number of repeating units. The LPS of the DPPE/LPS-5ru membrane, reorganize themselves as function of the hydration level (panel B) as shown by the density distributions along the membrane normal of the different components (panel C). In panel B, dark blue spheres represent Cl^−^, cyan spheres Na^+^, green spheres Ca^2+^ and water molecules are presented as blue spheres (top half of the water on the LPS leaflet was removed for clarity).

The inner leaflet initial configuration was generated with the CHARMM-GUI membrane builder.^25–30^ The outer leaflet was built by placing the LPSs in a 4*4 grid, with random rotation along the membrane normal and minimal spacing to avoid any collisions of atoms.

Water and ions are then added through VMD^31^ by using the Solvate and Autoionize plugins. NaCl 0.15 m was added to the system in addition to 80 Ca^2+^ counter ions needed toion 36^18^ for lipids and TIP3P for water molecules.^19^ MD simulations were performed neutralize the residual negative charges of the outer leaflet. NaCl ions were randomly placed in the water, while the Ca^2+^ ions were placed in the region corresponding to the inner and outer core of the LPS (1.5 nm to 4.0 nm relative to the center of the bilayer hydrophobic region).

Equilibration of the systems was obtained by 8 cycles of short (5 ns) alternating NPsT and NVT simulations. During the NPsT simulations, a gradually vanishing surface tension tension bias (starting at −0.400 N/m) was applied to compress the system. After these cycles, the systems were equilibrated for 200 ns under NPsT conditions. The membranes were assumed to be at equilibrium once the time average of the area per lipid changed less than 10%, as shown in Fig. S2.

The equilibrated structures were used to prepare the dehydrated systems. We prepared five dehydrated systems for each bilayer, denoted as 83%, 67%, 50%, 33% and 17%, with the percentage indicating the amount of remaining water in the outer leaflet, as described below. To prepare a system at a given hydration level (*x*%) we:

1. remove a number of water molecules on the outer leaflet that allowed us to keep exactly the *x*% of the molecules that were present from the hydrophobic center up to the last residue of the repeating units in the fully hydrated system.
2. left the water of the inner leaflet unchanged (that is a layer of 4 nm or more).
3. removed NaCl ions to keep a 0.15 m concentration on both sides.
4. run a 200 ns of simulation in NVT ensemble to equilibrate the system.

In all cases, the analyses, performed with VMD^31^ and MDAnalysis, ^32,33^ were based on the last 100 ns of each simulation. ACD ChemSketch^34^ was used to draw molecule skeletal structure.

### Diffusion coefficients

The diffusion coefficient of water was calculated from the integration of velocity autocorrelation function. ^35^ To this end, we run short (100 ps) simulations of each system, saving atomic velocities every 10 fs. To calculate the water diffusivity in different regions (defined in Fig. 4), we selected only the water molecules within 1 nm, along the membrane normal, from the center of each region. To reduce numerical noise, the autocorrelation functions of the velocity were integrated up to 2 ps.

## Results

The exact structure of the outer membrane of Gram-negative bacteria is highly variable and therefore we needed to select only some representative cases to study the effect of dehydration on their structure and dynamics.

DPPE/LPS-5ru membranes were simulated with different amount of water on the LPS side, as shown in Fig. 1. The fully hydrated one (100%) contains the DPPE/LPS-5ru membrane with enough water to saturate the full structure plus an additional 6 nm layer of water to emulate bulk water. A set of 5 dehydrated membranes (83% to 17%) was generated from the fully hydrated one by removing part of the water in the LPS region and exposing it to the vacuum. As detailed ion the methodology, the different hydration levels were labelled based on the percentage of water left in the LPS region.

As expected the exposure of the LPS chains to air, causes them to collapse (see Fig. 1B), but the effect is mostly limited to the repeating units, while the other regions are barely affected except at the highest dehydration level (17%). At the lowest hydration levels the water loses its cohesion as small amounts of water remain in the repeating unit region rather than the outer core. Of note, the evaporation on the LPS is negligible, and in general no flux of water molecule leaving the LPS is observed.

The density distributions along the membrane normal of the various components, show the LPS collapse even more clearly (Fig. 1C). The peak-to-peak distance from the first to the last repeating unit drops from 4.5 nm, in the fully hydrated membrane, to 3 nm in the 17% hydrated system. At the same time dehydration causes the outer core, inner core and the lipid A head group to shift about 0.5 nm toward the hydrophobic center of the membrane. In contrast, there is no significant change on the DPPE leaflet, where the peak of the phosphorous group of the DPPE is about 1.6 nm below the membrane hydrophobic center at all hydration levels. There is also a noticeable change in the water distribution in the LPS region (Fig. 1C) as the density of water in the repeating-unit region decreases from 500 kg m^−3^ to less than 120 kg m^−3^ in the 17% hydrated one. However, the density of water near the inner core region remains around 500 kg m^−3^ at all hydration levels, likely due to the high ionic concentration.

The observed structural changes match the finding of previous works^2,11,36,37^ that have indicated the importance of repeating units in helping bacteria survive dehydration by maintaining a hydrated layer close to the cells that slows the surface evaporation rate. Our simulations show how, by forming a denser shell, the repeating units region traps the water close to the inner regions, which is known to be essential for the biological functions of membranes^38–43^ and, at the same time, reduces the evaporation rate. The latter is not only a steric effect, related to the tortuosity of the path, but also caused by the attractive interactions between the water and the repeating unit, as shown at 17% hydration where a fraction of water molecules is stabilized by the interaction in the region between 5 nm and 7.5 nm.

The simulations show that the structural adjustments to dehydration go beyond a simple change in the density distributions, but affect also the overall shape of the LPS molecules, as can be seen in Fig. 2. This change can be observed by monitoring the effect of the dehydration on z-axis distance (*D*_*RU*_) and the tilt angle (*θ*_*RU*_). These quantities, which measure the average distance from the membrane surface (*D_RU_*, see Fig. 2A) and tilt from the membrane normal (*θ*_*RU*_, see Fig. 2D), show the change in LPSs’ configurations.

**Figure 2:**
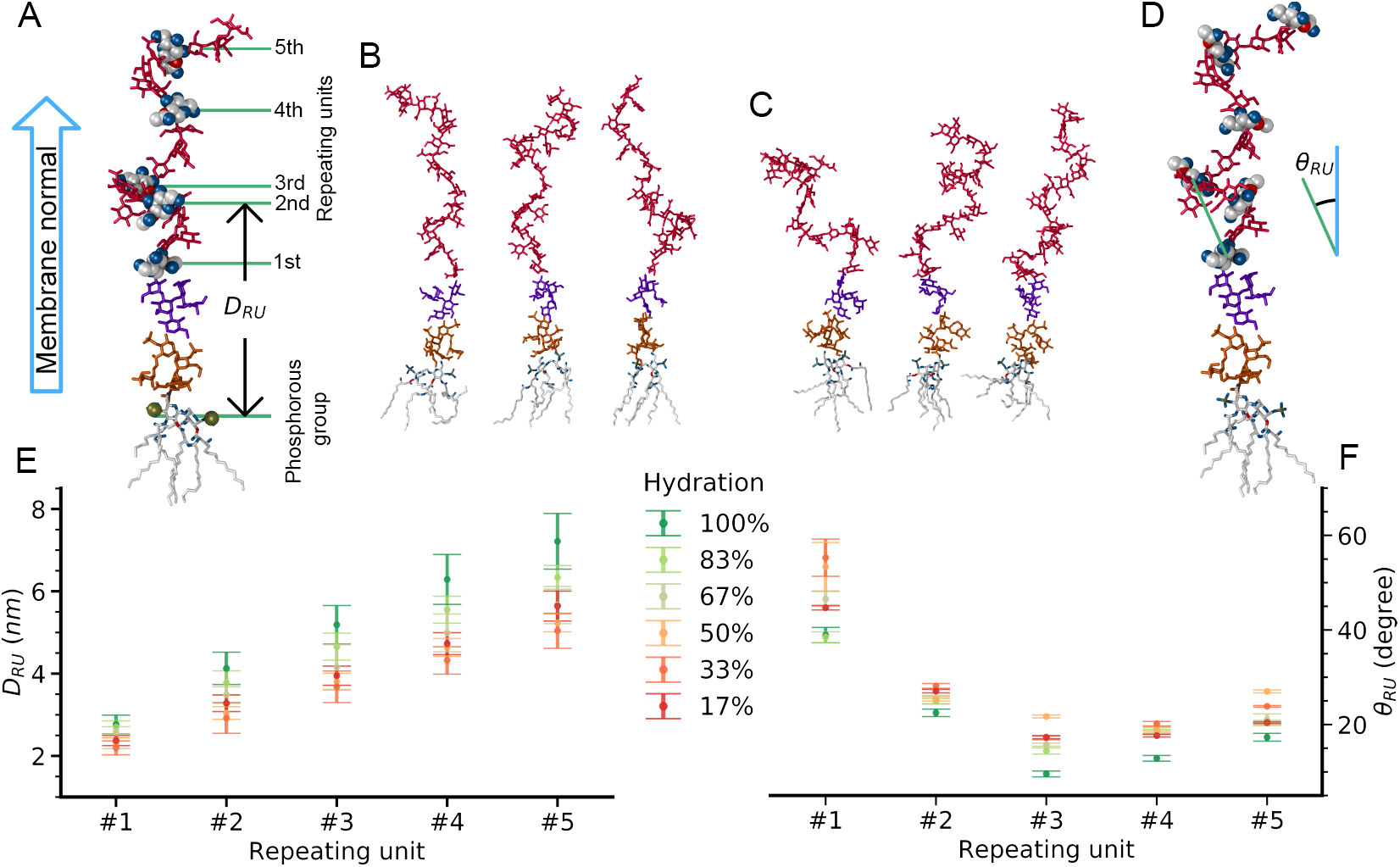
Structure of individual LPSs in the DPPE/LPS-5ru membrane. LPSs’ conformations were characterized by z-axis distance (*D*_*RU*_, panel A and E) and tilt angle (*θ*_*RU*_, panel D and F), with error bars equal to one standard deviation. Selected LPSs’ conformations at 100% and 17% hydration are shown in panel B and panel C, respectively.

*D*_*RU*_ (Figs. 2E) show that the compression along the membrane normal is roughly linearly correlated with the dehydration up to 50% but at higher desiccation the rest of the LPS are responsible for most of the outer membrane changes, while the thickness of repeating units remains roughly unchanged. *θ*_*RU*_ follows a similar trend with increasing angles at lower water contents. Interestingly, in both cases at the lowest water content, the evaporation of water from the core region to the repeating units causes a partial restoration of the more extended structure typical of higher hydration levels (67%).

To investigate the correlation between the effects described above and the presence of re-peating units in the LPS molecules, we compared the results, under similar conditions, of an identical membrane except for the lack of repeating units (DPPE/LPS-0ru). The structure of one of the LPS is shwon in Fig. 3C, together with the structures and corresponding mass distributions of the DPPE/LPS-0ru membranes in at different hydration levels (Figs. 3A and 3B).

**Figure 3:**
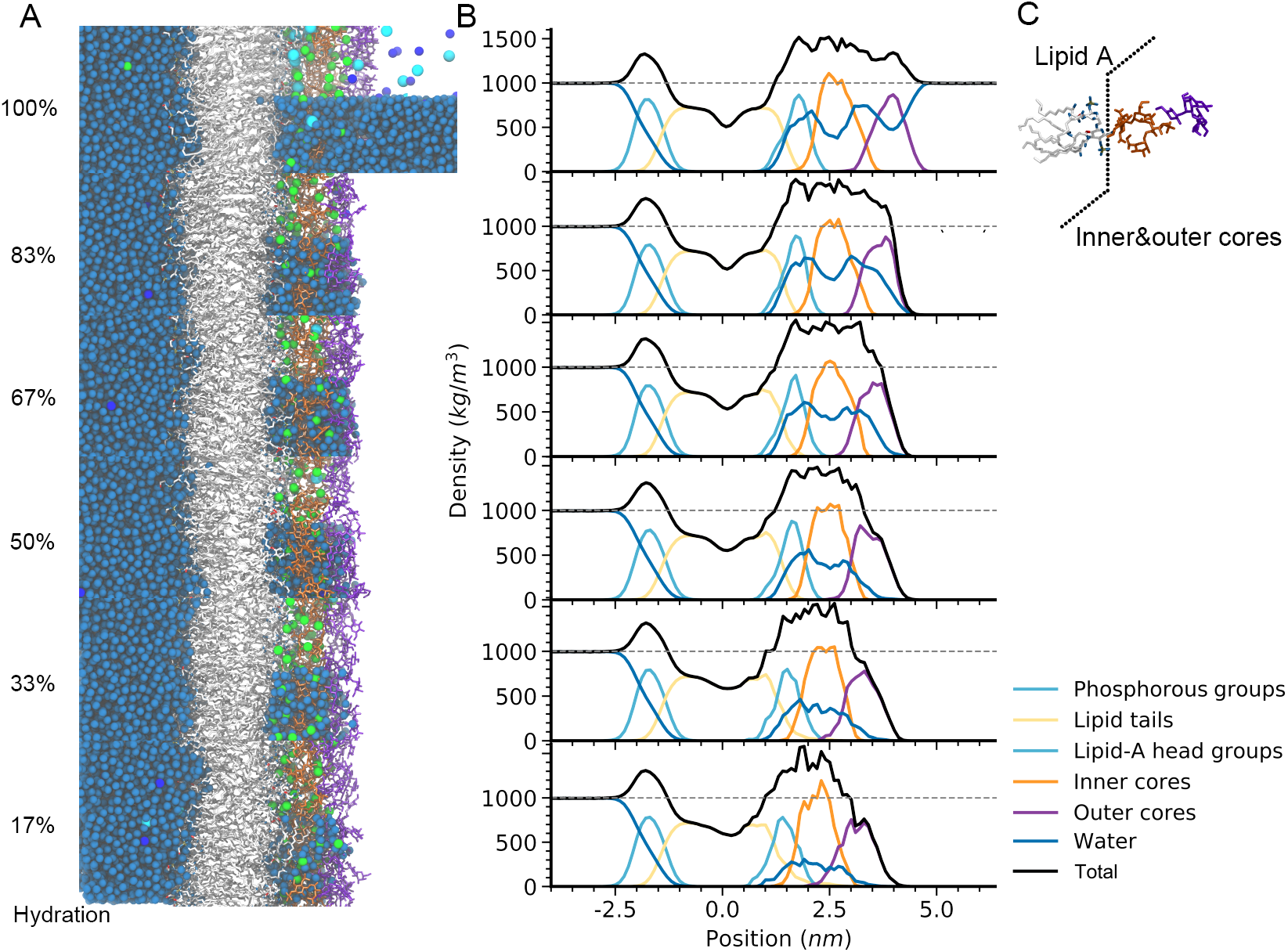
Structure and mass distribution of the DPPE/LPS-0ru membrane. LPS structure is shown in panel C and contains lipid A, inner core, outer core but no repeating units. The effect of hydration can be seen in the structure (panel B; see Fig. 1 for details) and the density distribution (panel C).

By comparing these results with Fig. 1, we can see how the outer core (based on the position of the density distribution peak) shifts of about 0.5 nm toward the membrane as it did for DPPE/LPS-5r system. However in contrast with the previous case, the water depletion is proportional to the level of dehydration and the inner layer of water is not preserved, as shown by a water density (200 kg m^−3^ at 17% hydration), which is about halved compared to the system with 5 repeating units. While a minimal water-depleted stratum is formed, that can potentially slow the evaporation, it is markedly thinner than the one observed before, which can be linked to a reduced water retention capability for LPS lacking repeating units.

## Discussion

### Water diffusivity

Based on the previous results, it is reasonable to expect that the condensed layer of repeating units may shield the cell from the mass exchange between cytosol and the environment, especially to prevent the loss of water during desiccation. To validate this hypothesis, we computed the water diffusivity in each region of the outer membranes for both systems at all the hydration levels. The relative change in diffusivity compared to self-diffusivity of water in bulk is reported in Fig. 4.

**Figure 4:**
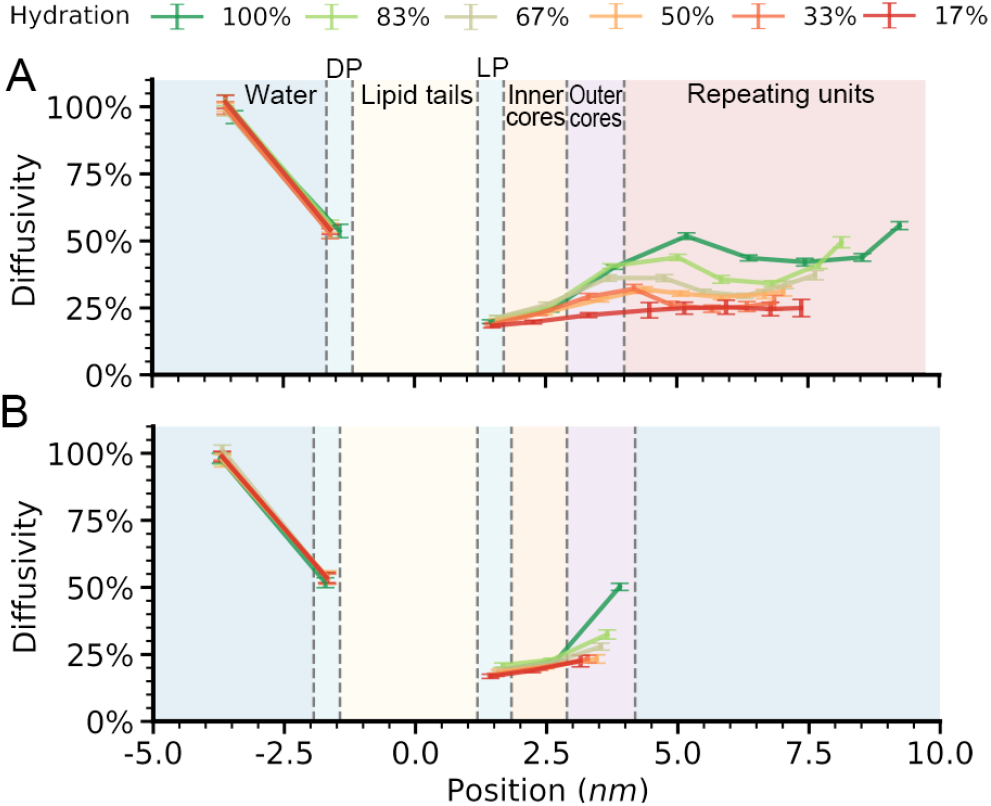
Relative diffusivity of water in the (A)DPPE/LPS-5ru and (B) DPPE/LPS-0ru membranes as a function of the hydration levels. The value shows the water self-diffusion in different regions relative to bulk water, with error bar corresponding to one standard deviation. DP stands for the DPPE phosphorous group and LP stands for the lipid A phosphorous group.

The result show a direct correlation between the hydration level and the water diffusivity in all the regions. The water diffusivity drops form about 50% in the most external region of each layer (repeating units for the DPPE/LPS-5ru and outer core for DPPE/LPS-0ru) to approximately 25%. The similarity between the two systems, leads us to conclude that the dense layer that is formed during dehydration acts similarly, independently of the presence of repeating units, but the increased thickness of the low diffusion region, significantly reduces the water loss in desiccated environment when the repeating units are present. Interestingly, the repeating units halve the diffusivity of water even in the fully hydrated system, and we expect a much larger effect for larger molecules. Finally, we expect the reduction in diffusivity to larger than the one predicted by the simulations. The TIP3P model used for water in these simulations is known to overestimation the diffusivity under 323 K (50 °C).^44,45^ For example, we measured a bulk diffusion coefficient of 4.65 × 10^−9^ m^2^ s^−1^ at 310 K (37 °C), instead of the expected value of 3.04 × 10^−9^ m^2^ s^−1^.^46^ This discrepancy is greater when water diffusivity is reduced (*e.g.*, at lower temperatures), ^44^ leading us to believe that while the overall trends were corecly captured by MD, the effective reduction is larger.

### Atomic mobility

Beside the diffusivity of water molecules, the other factor that strongly characterizes the biological activity of the membranes is its rigidity, as it affects many critical processes, like passive transport and transmembrane protein mobility. To quantify the membrane molecules rigidity, we measured the average root mean square fluctuations (RMSF) of the membranes atoms, that is 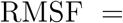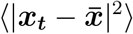, where ***x***_***t***_ is the position of an specific atom at time *t*, 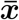 average position and angular brackets denote ensemble average. The average RMSF of each region in the LPS membranes at different hydration levels is shown in Fig. 5.

**Figure 5:**
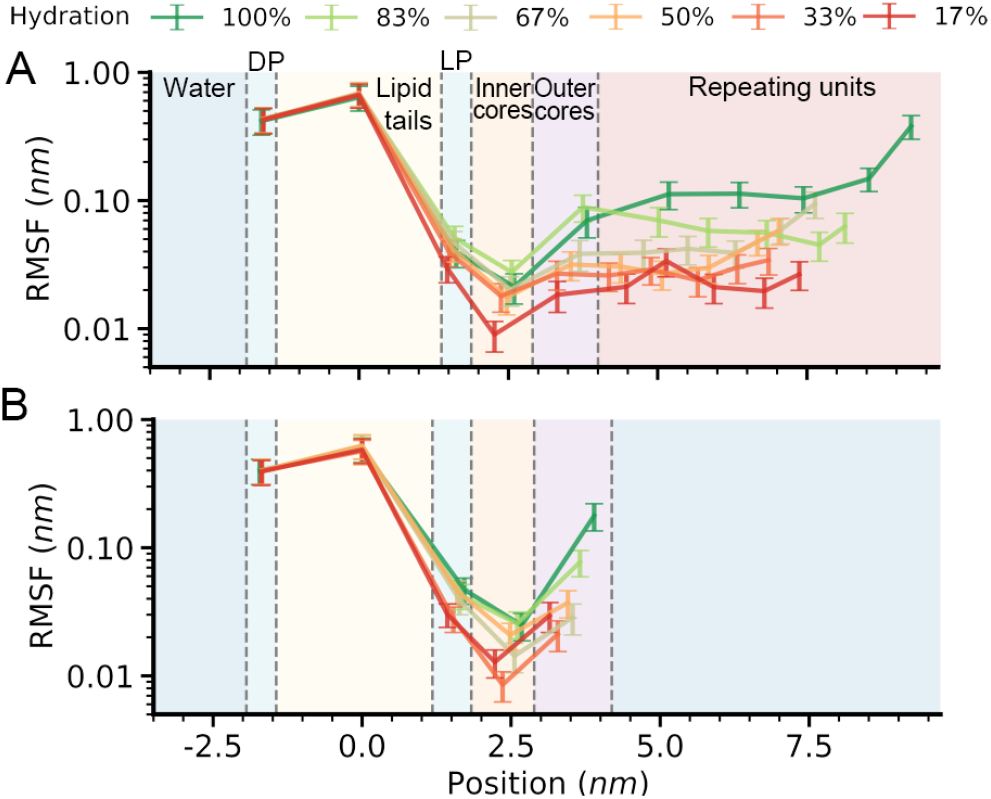
Atomic mobility measured as root mean square fluctuations (RMSF) of the positions in the (A) DPPE/LPS-5ru and DPPE/LPS-0ru membranes at different hydration levels. Error bars equal to one standard deviation.

The results for the fully hydrated membrane show expected features, for example an increased mobility of the lipid tails in the membrane center or the outermost part of the LPS (repeating units or outer core for DPPE/LPS5ru and DPPE/LPS-0ru, respectively). While there is a generally positive correlations between the RMSF and the membranes’ density (Fig. 1C) and therefore, would be intuitive to associate the increased mobility with a local decrease in atomic density, that is not always the case. For example, when comparing the DPPE/LPS-5ru system at 83% and 100% hydration level, we observe a decrease in the RMSF in the repeating units region, despite no significant difference in the region’s density. Of note, the comparison between the two fully hydrated systems, shows that the presence of the repeating units does not significantly affect the mobility of the atoms of the inner leaflet, the hydrophobic region or even the surface of the outer leaflet (core and LP regions).

The desiccation process results in general in a more rigid outer layer with a loss of almost one order of magnitude in atomic mobility. The presence of the repeating units however causes this reduction to be more uniform across the repeating unit region, independently of the water distribution, resulting in an overall more rigid shell compared to the LPS lacking repeating units.

### Hydrophobic region

Another common atomistic descriptor of the membrane characteristics is the hydrophobic thickness, which has been shown to be related to several phenomena like passive transport^47^ and membrane-protein binding.^48^ In this work, the hydrophobic thickness is defined as the average distance between the C2 (for acyl chain number 6 and 8) and C4 (for other acyl chains) carbon atoms on lipid A and the C2 carbon atoms of DPPE (see Fig. S1), and the results for the two systems at different hydration layer are shown in Fig. 6.

**Figure 6:**
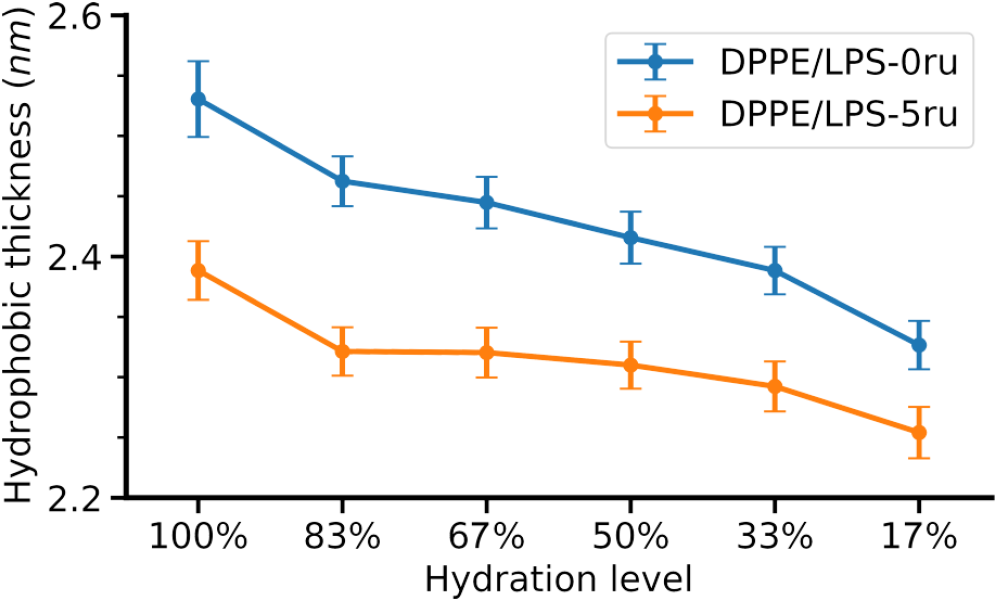
Hydrophobic thickness of the DPPE/LPS-5ru and DPPE/LPS-0ru membranes at different hydration level. Error bars equal to one standard deviation.

For the fully hydrated systems, the DPPE/LPS-0ru and DPPE/LPS-5ru membranes differ of 0.14 nm, in perfect agreement with published results of 0.28 nm for a double LPS membrane^16^ (namely LPS0 and LPS5 in the cited work). At decreasing hydration levels, the hydrophobic thickness decreases, which corresponds to a increased membrane permeability, all other factors being the same. This increase is a consequence of the low compressibility of the membrane, so a reduction in thickness corresponds to an increase in are per lipid. The repeating unit limit the increased permeability, by increasing the attractive intermolecular LPS interactions, which in turn curbs the increase in area per lipid.

The change in hydrophobic thickness, is the result of a reorganization in the lipid tails, as can be shown by the evolution of the relative orientation of the carbon atoms in the acyl chains. The lipid tail conformation was characterized by *θ_CC_*, which is the angle between the vector defined by the position of two consecutive carbon atoms and the membrane normal.

The results for representative acyl chains (chain 1 of lipid A, see Fig. S1 in the Supplementary Material for chain indexing) are shown in Fig. 7. Despite the contraction of the hydrophobic layer, we observe no relevant difference in the lipid tail organization except at hydration levels of 33% or lower for the DPPE/LPS-5ru membrane, while the desiccation starts affecting the DPPE/LPS-0ru membrane earlier. This effect is generally valid even though interestingly the response to the hydration is somewhat varies for each chain (see the full set of results in Figs. S3 and S4 in the Supplementary Material).

**Figure 7:**
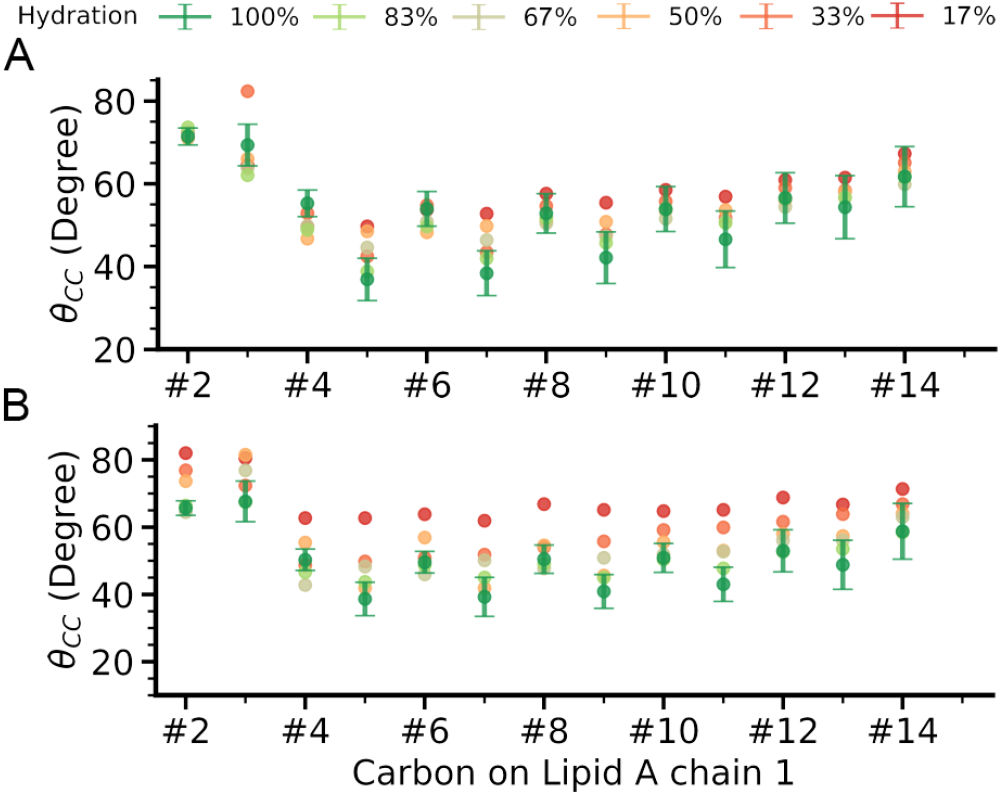
C–C tilt angle of chain 1 of lipid A (see Fig. S1) in the (A) DPPE/LPS-5ru and (B) DPPE/LPS-0ru membranes at different hydration. Error bars, equal to one standard deviation, are similar for all the hydration level and therefore only the 100% is shown for clarity.

The bending of the lipid tails generally increases at lower water contents, which corresponds to the reduction in hydrophobic thickness, but the relative change is larger for the system without repeating units, as shown in Fig. 8, which shows the overall change for all the acyl carbons relative to the fully hydrated system.

**Figure 8:**
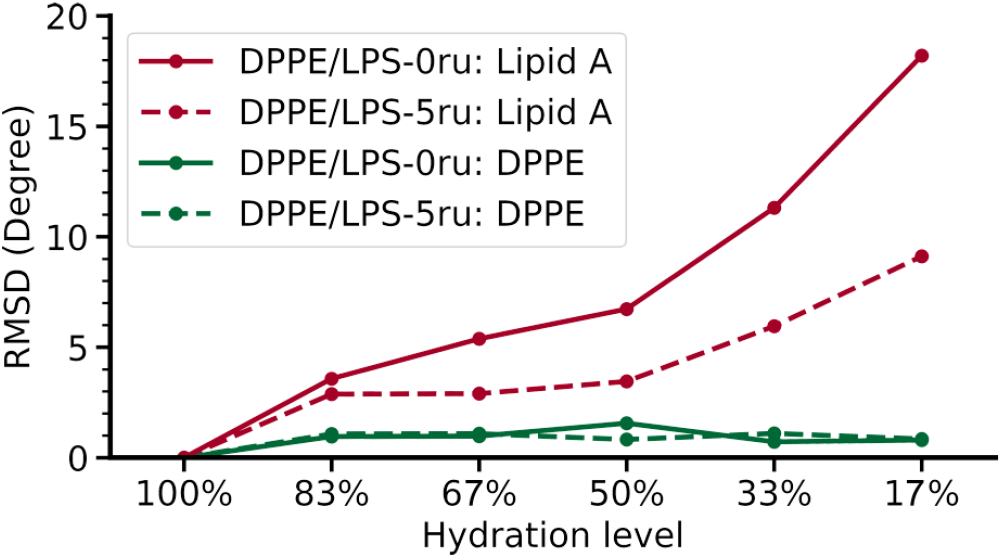
Root mean square difference of *θ*_*CC*_ for all the chains relative to the fully hydrated system.

The data show that at increased desiccation, the acyl chains in the system without repeating units have much higher tendency to lay more perpendicular to the membrane normal. As the acyl chains of the DPPE is not significantly affected by dehydration process, this effect can only be ascribed to the presence (or absence) of repeating units. This marked change in structure for DPPE/LPS-0ru is likely to further affect lipid packing, lipid lateral diffusion,^49^ membrane surface tension,^50^ permeation^51^ and membrane protein binding.^52^

## Conclusion

In this work, we investigated the short term response of Gram-negative bacteria to desiccation, specifically the role that the outer membrane has in resisting dehydration.^1,2,9^ By using molecular dynamics simulations, we have analyzed the effect of different water content on *E. coli* LPS model membranes and the change of the membrane properties for different composition of the LPS molecules.

Our results, which are in agreement with previous published work,^14–16,53^ show that the presence of even a small number of terminal repeating units on the LPSs has profound effects in the dehydration response. The repeating units are responsible for two major effects: creating a layer of polysaccharides that reduces water loss and maintains the general characteristics and functionality of the membrane. The presence of the polysaccharide layer allows the persistence of a small water layer close to the membrane hydrophobic region at all but the most extreme dehydration levels and generates a more rigid shell as shown by the atomic mobility, water diffusivity and density distributions. This protective outer layer also indirectly affects the hydrophobic region (*i.e.*, hydrophobic thickness, and acyl chain orientation) of the outer membrane by preserving its structure, which is critical for maintain the functionality of the membrane in desiccated environment.

All these structural effects are general and thermodynamically driven, resulting in a rapid and repeatable response of bacteria to the desiccation stress that occurs before the slower biosynthetic and gene expression adjustments can take effect. Therefore, we expect that the application of this understanding and the approach shown here can be beneficial for future studies on the role of bacterial outer membrane in surviving environmental stress, with applications that span the developing of antimicrobials to more dehydration resistant bacteria for agriculture.

## Acknowledgement

This work was funded by the University of Michigan College of Engineerings Blue Sky Initiative.

## Supporting Information Available

Additional information about molecular structure and MD simulations.

## Notes

### Competing Interest Statement

The authors have declared no competing interest.

