## Supplementary Information for "Molecular Dynamics Study of Dehydrated Lipopolysaccharide Membrane"

### Supporting Information
### Molecular Dynamics Study of Dehydrated Lipopolysaccharide Membrane

Changjiang Liu,† Paolo Elvati,† and Angela Violi*,†,‡,¶

†*Biophysics Program, University of Michigan, Ann Arbor, MI 48109-2125, USA*

‡*Department of Mechanical Engineering, University of Michigan, Ann Arbor, MI 48109-2125, USA*

¶*Department of Chemical Engineering, University of Michigan, Ann Arbor, MI 48109-2125, USA*

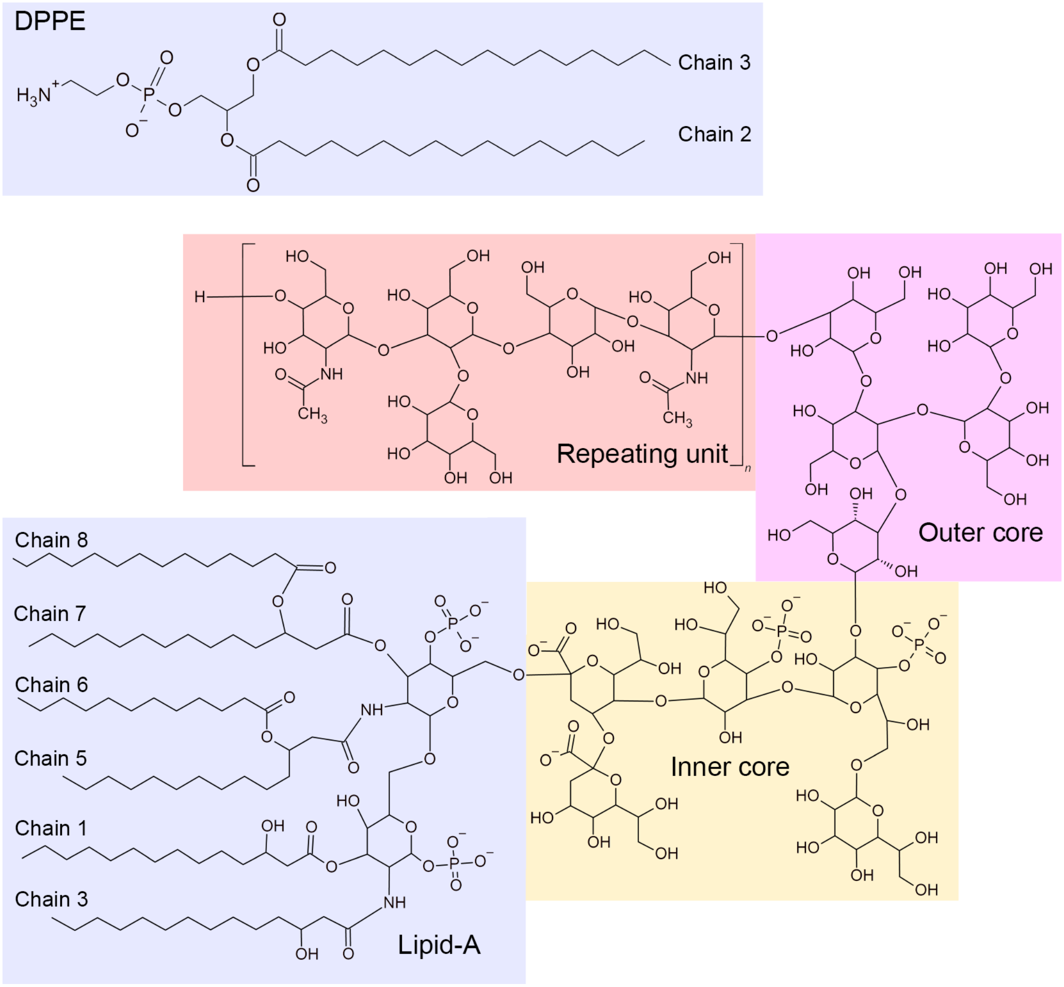


Figure S1: Molecular structure of the DPPE and LPS in this work. Fatty acid chains are indexed as indicated on the plot.

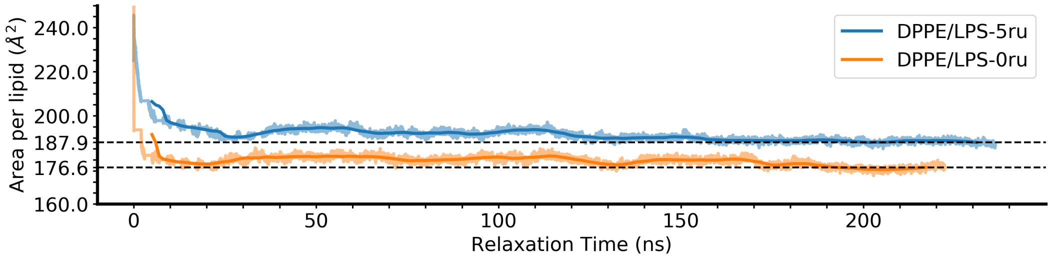


Figure S2: Relaxation process of the DPPE/LPS-5ru and DPPE/LPS-0ru membrane. The solid line is the running average over a 10 ns window.

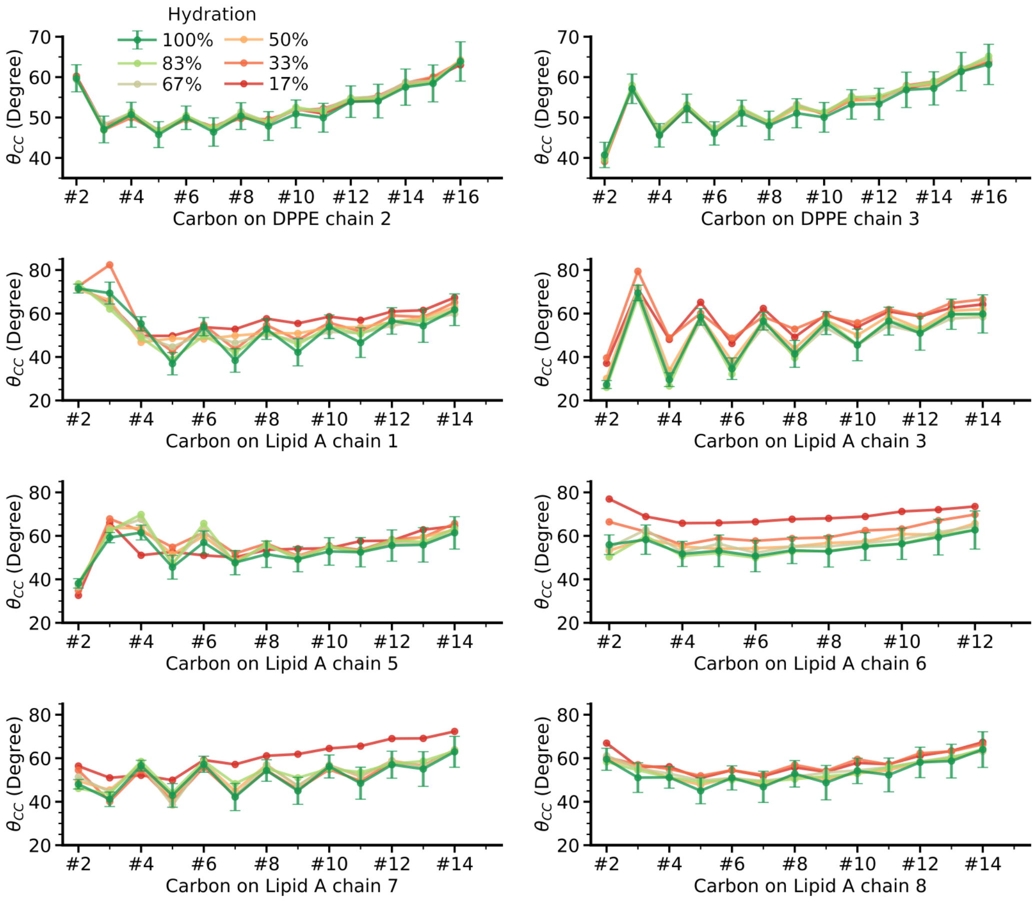


Figure S3: C-C bond tilt angle for each acyl chains in the DPPE LPS-5ru membrane. Refer to Figure S1 for the index of chains. Error bars represent one standard deviation, and are shown only for the hydration level 100% examples for clarity.

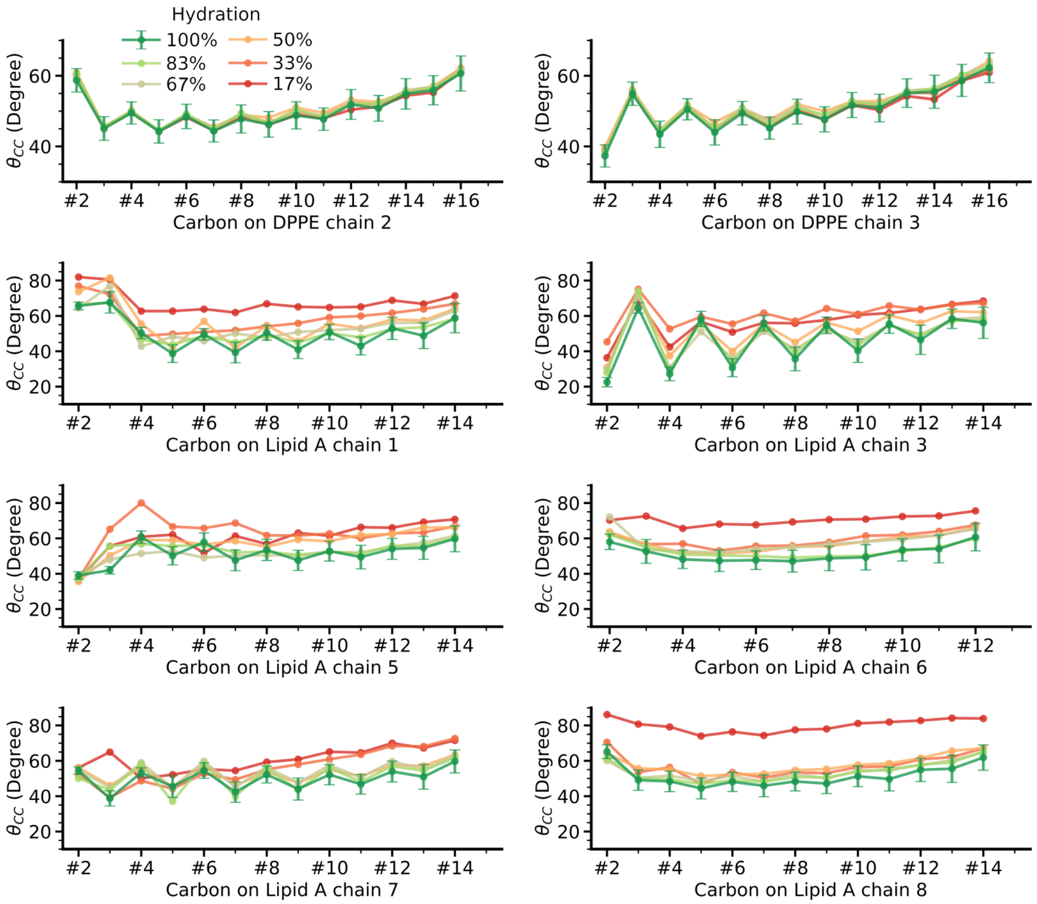


Figure S4: C-C bond tilt angle for each acyl chains in the DPPE LPS-0ru membrane. Refer to Figure S1 for the index of chains. Error bars represent one standard deviation, and are shown only for the hydration level 100% examples for clarity.
